# Ketone body 3-hydroxybutyrate alleviates CCl4-induced liver fibrosis in mice through regulation of mitochondrial fission and reduction of oxidative stress

**DOI:** 10.1101/2024.01.27.577553

**Authors:** Yudian Zhang, Xinyi Liu, Yifan Wang, Mengyuan Liu, Ziyi Guo, Jinbo Zhang, Fuqing Wu, Guo-Qiang Chen

**Author notes:** **CORRESPONDING AUTHOR :** Guo-Qiang Chen,l School of Life Sciences, Tsinghua University, Beijing 100084, China, Yudian Zhang, Department of Cell Biology, School of Basic Medical Sciences, Capital Medical University, Beijing, 100069 P. R. China. These authors contributed equally (Yudian Zhang, Xinyi Liu).

## Abstract

3-Hydroxybutyrate (3HB) is an important metabolite and regulatory molecule produced in liver. Previous studies have shown that 3HB could be beneficial to many diseases, including brain diseases, diabetes, and most importantly, inflammation and liver related injury. Therefore, the effect of 3HB on liver fibrosis, one key step of liver diseases which proved to be reversible, is urgent to explore. In this study, the CCl4-induced mouse model of liver fibrosis has been successfully constructed and treated by 3HB. The results demonstrate that 3HB could alleviate CCl4-induced liver injury and inflammation in mice, decrease the accumulation of collagen, the expression of pro-fibrotic genes as well as inflammatory factors, and finally the degree of liver fibrosis. The transcriptome data recovers that the anti-fibrotic effect of 3HB might be exerted through several ways, such as regulating mitochondrial function, reducing oxidative stress and p53 signaling pathways, proposing a safe and relatively fast possibility for the treatment of liver fibrosis.

## INTRODUCTION

Chronic liver diseases are a major global health burden and account for approximately 2 million deaths per year worldwide[1]. As the middle stage of stage of liver diseases, Liver fibrosis is a key factor for liver disease outcome and risk of hepatocellular carcinoma (HCC)[2]. Liver fibrosis occurs when accumulation of extracellular matrix (ECM) proteins, mostly crosslinked collagens type I and type III, which replace damaged normal tissue and form a fibrous scar[3]. The main causes of liver fibrosis include chronic hepatitis virus infection, alcohol abuse, and nonalcoholic steatohepatitis (NASH) in modern society[4]. The accumulation of ECM proteins distorts the hepatic architecture and subsequently developed into cirrhosis, which results in hepatic insufficiency and portal hypertension[5]. In 1990s, researchers confirmed that even advanced liver fibrosis could be reversible, which attracted much attention in developing various antifibrotic therapies to suppress the progression of series of liver diseases at the early stage[6]. The common antifibrotic therapies nowadays include inhibition of hepatocyte apoptosis; reduction of oxidative stress; inhibition of HSC activation and reduction of fibrotic scar evolution and contractility[7]. However, there is still not yet standard treatment for liver fibrosis. The current selected anti-fibrotic candidate agents are either too slow or too infrequent to avoid life-threatening complications in late-stage disease particularly, with some ineffective when move to clinical trials[8]. Therefore, effective, safe and hopefully more rapid drugs are urgent for this market.

3-hydroxybutyrate (3HB) is a very important metabolite and regulatory molecule which occurs in bacteria and animals[9]. As the most abundant ketone body in mammals, 3HB is able to provide energy instead of carbohydrate under extreme conditions, such as fasting or prolonged exercise[10]. Under these circumstances, fatty acids are mobilized from adipocytes and transported to the liver for conversion to ketone bodies, and then be distributed via blood circulation to metabolically active tissues to be metabolized into acetyl-CoA and eventually ATP[11]. 3HB is also regarded as a therapeutic agent with beneficial effect for many diseases[12]. For instance, 3HB or exogenous 3HB supplementation can affect brain diseases, such as epilepsy, multiple sclerosis (MS), stroke, depressive disorder, and schizophrenia [13]. More closely related to this project, 3HB or ketogenic diet attenuates caspase-1 activation and IL-1β secretion in mouse models of NLRP3-mediated diseases such as Muckle-Wells syndrome, familial cold autoinflammatory syndrome and urate crystal-induced peritonitis[14]. Also, deficiency of 3HB dehydrogenase (BDH1) in mice causes low ketone body levels and fatty liver during fasting, which is the former step of liver fibrosis[15]. According to our previous study, 1,3-BDO, a precursor substance of 3HB, could reduce liver and kidney injury caused by type 2 diabetes, and improve liver function[16]. It was also reported that 3HB protects from alcohol-induced liver injury through a Hcar2-cAMP dependent pathway [17]. Based on the above evidence, it is reasonable to suggest a hypothesis that 3HB could also alleviate or even reverse liver fibrosis to some extent.

In this study, 3HB was administrated to mice with CCl_4_-induced liver fibrosis for the first time as a therapeutic agent. The results showed that 3HB could significantly improve liver function, inhibit the expression of liver fibrosis markers and pro-inflammatory factors in this animal model, therefore ameliorate liver fibrosis in terms of both phenotype and genetics.

## RESULTS

### Different doses of 3HB all exhibit attenuation of CCl_4_-induced liver fibrosis in mice

Previously, we found that 3HB had a protective effect on the liver of mice with type 2 diabetes[16]. CCl_4_ directly causes liver cell injury, necrosis and inflammation, and is widely used in the establishment of animal models of liver fibrosis and cirrhosis[18]. In order to further explore whether 3HB has a therapeutic effect on other more serious progressive liver diseases, we induced liver fibrosis in mice by intraperitoneal injection of CCl_4_ for a continuous 4 weeks, while administering different concentrations of 3HB, to investigate whether 3HB could have a therapeutic effect on liver fibrosis. H&E, Masson and Sirius red staining were used for the analysis of liver injury and fibrosis.

H&E staining results indicate that the liver structure of mice treated with olive oil is normal. However, after 4 weeks of CCl4 administration, necrosis and inflammatory cell infiltration occurred in the central lobule of the liver in the mice treated with CCl_4_. These pathological changes were inhibited by different doses of 3HB (Figure 1A), suggesting that 3HB can alleviate the liver damage induced by CCl_4_ in mice.

**Fig 1.**
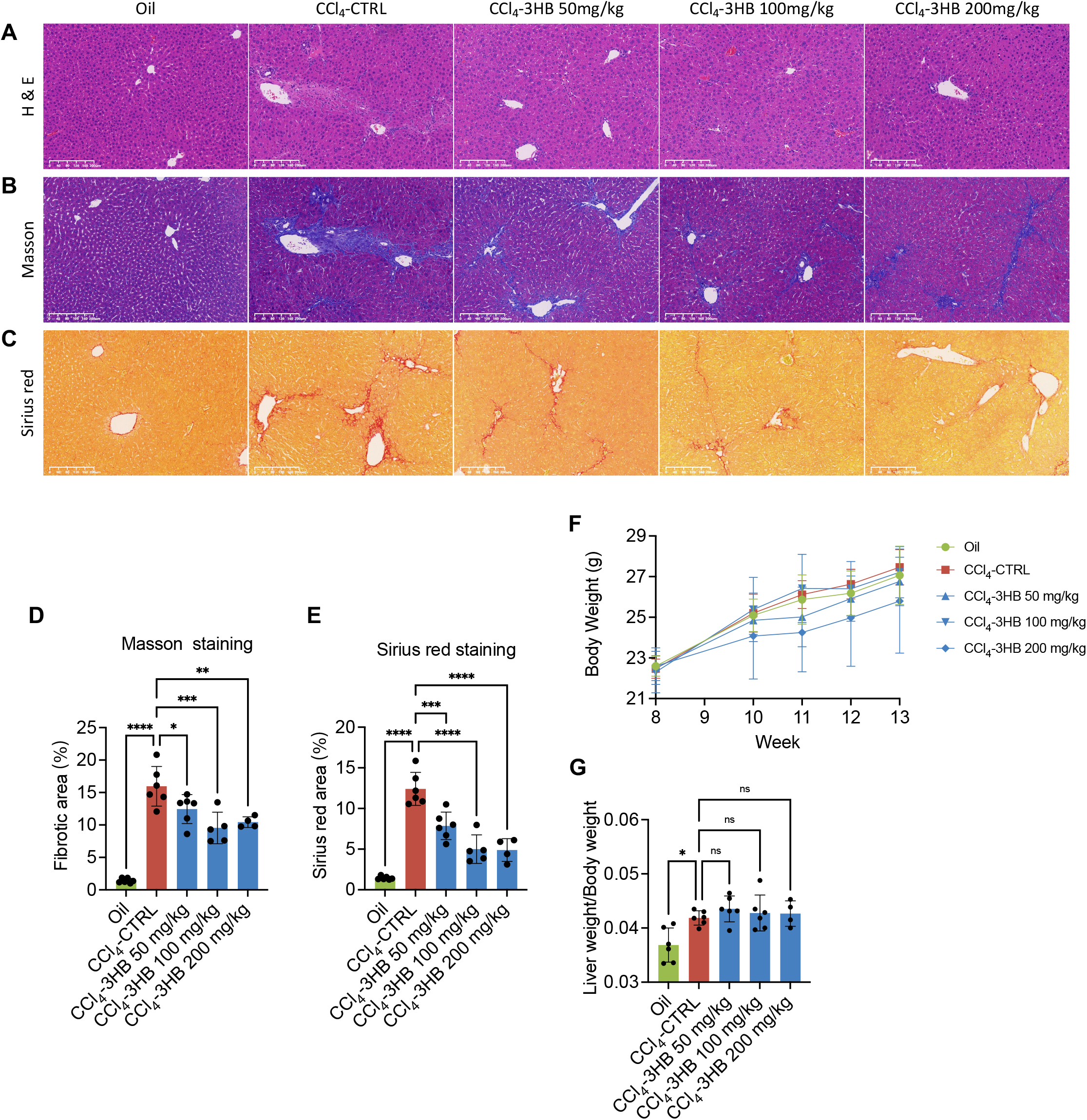
Different doses of 3HB all exhibit attenuation of CCl_4_-induced liver fibrosis in mice. A-C. Representative H&E, Masson and Sirius red staining images of livers after 4-week CCl_4_ treatment; scale bars, 200 μm. D-E. Quantification of fibrosis by relative lesion size. F. Body weight for 6 consecutive weeks. G. The ratio of liver weight/body weight at the end of the experiment. Date is presented as meansL±LS.E.M. (nL=L4-6). *p□<□0.05 and **p□< □0.01, ***p□< □0.001, ****p□< □0.0001, compared with CCl_4_-CTRL group.

Masson’s staining revealed a significant proliferation of the connective tissue in the livers of mice treated with CCl_4_ compared to those treated with olive oil, indicating the successful establishment of a liver fibrosis model. Treatment with different doses of 3HB significantly reduced the accumulation of collagen fibers in the liver (Figure 1B & 1D), demonstrating a significant protective effect of 3HB on liver fibrosis.

In Sirius red staining, collagen was significantly accumulated in the livers of mice treated with CCl_4_, but various doses of 3HB noticeably reduced this accumulation (Figure 1C & 1E). The aforementioned staining results suggest that 50 mg/kg, 100 mg/kg, and 200 mg/kg of 3HB can all alleviate the liver tissue damage and the degree of liver fibrosis in mice treated with CCl_4_, without negative impact on the body weight or relative liver weight of the mice (Figure 1F-1G). Among these, the dose of 100 mg/kg of 3HB can be selected as the optimal dose for subsequent experiments.

### Different doses of 3HB can improve the liver function of mice with liver fibrosis

To further investigate the protective effect of 3HB on the livers of mice with liver fibrosis, we measured the levels of alanine aminotransferase (ALT), aspartate aminotransferase (AST), and total bilirubin (TBIL) in serum. Compared with the olive oil treatment group, liver function in the CCl_4_ treatment group was impaired. However, different doses of 3HB significantly reduced the levels of ALT, AST, and TBIL in the serum of mice with liver fibrosis (Figure 2A-2C), reversing the liver function damage caused by CCl_4_.

**Fig 2.**
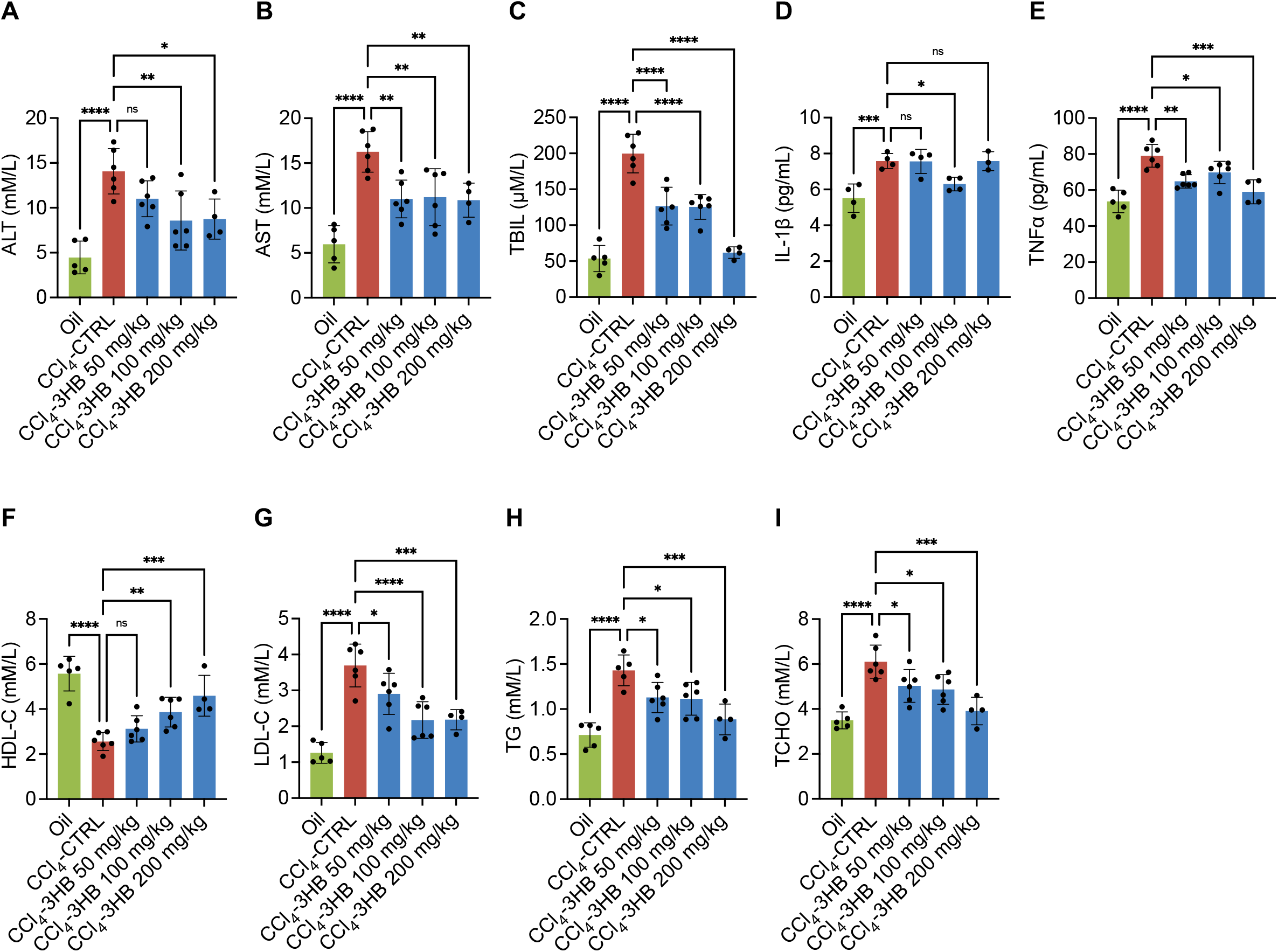
Different doses of 3HB can improve the liver function of mice with liver fibrosis. A. Serum ALT level. B. Serum AST level. C. Serum TBIL level. D. Serum IL-1β level. E. Serum TNFα level. F. Serum HDL-C level. G. Serum LDL-C level. H. Serum TG level. I. Serum TCHO level. Date is presented as meansL±LS.E.M. (n□= □4-6). *p□< □0.05 and **p□< □0.01, ***p□< □0.001, ****p□< □0.0001, compared with CCl_4_-CTRL group.

In addition, different doses of 3HB could also ameliorate the dyslipidemia in mice with liver fibrosis, significantly increasing the level of high-density lipoprotein cholesterol (HDL-C) and reducing the levels of low-density lipoprotein cholesterol (LDL-C), total triglycerides (TG), and total cholesterol (TCHO), suggesting that 3HB may act against dyslipidemia caused by liver fibrosis (Figure 2F-2I). Treatment with 3HB also significantly reduced the levels of inflammatory factors, IL-1β and TNFα, in the serum of mice with liver fibrosis (Figure 2D-2E), indicating that the therapeutic effect of 3HB on liver fibrosis may derive from its anti-inflammatory function.

### 3HB inhibits the expression of liver fibrosis markers

To further confirm the therapeutic effect and mechanism of 3HB on liver fibrosis, we administered a dose of 3HB 100 mg/kg via gavage in mice with CCl_4_-induced liver fibrosis. H&E staining results indicated that 100 mg/kg 3HB significantly reduced hepatic necrosis, lessened the infiltration of inflammatory cells, and decreased hepatocellular steatosis in mice with liver fibrosis (Figure 3A). Masson staining results suggested that 100 mg/kg 3HB notably suppressed the proliferation of connective tissue in the liver and reduced collagen accumulation (Figure 3B & 3E). Sirius red staining results also demonstrated that 100 mg/kg 3HB could inhibit collagen accumulation in the liver (Figure 3C & 3F).

**Fig 3.**
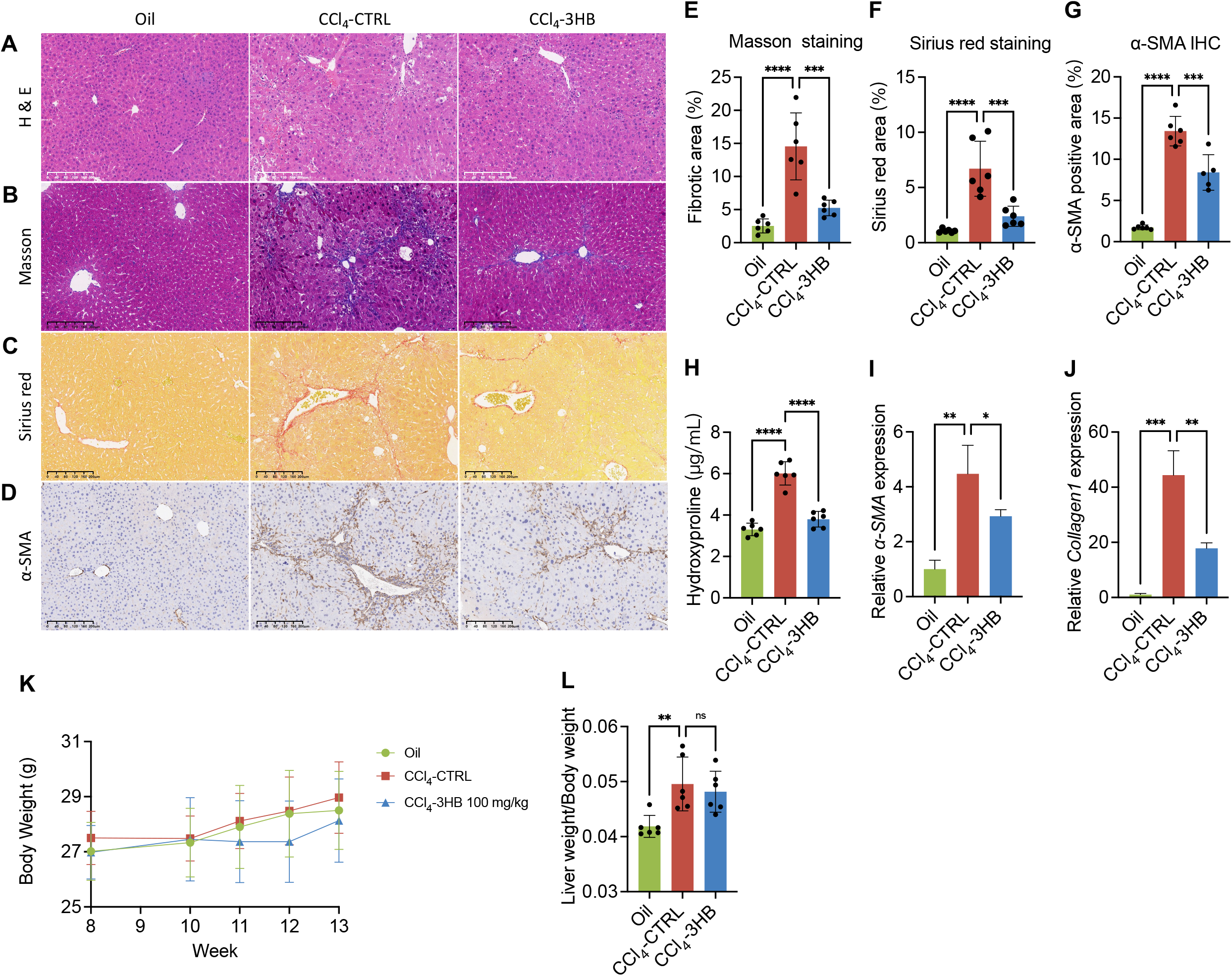
3HB inhibits the expression of liver fibrosis markers. A-D. Representative H&E, Masson, Sirius red staining and α-SMA immunohistochemical staining images of livers after 4-week CCl_4_ treatment; scale bars, 200 μm. E-F. Quantification of fibrosis by relative lesion size. G. Quantification of α-SMA positive area. H. Serum hydroxyproline level. I. The mRNA expression levels of α-SMAin liver. J. The mRNA expression levels of collagen 1 in liver. K. Body weight for 6 consecutive weeks. L. The ratio of liver weight/body weight at the end of the experiment. Date is presented as means□±□S.E.M. (n□=L6). *p□< □0.05 and **p□< □0.01, ***p□< □0.001, ****p□< □0.0001, compared with CCl_4_-CTRL group.

We also examined the expression levels of fibrosis markers in the liver, and found that the expression of α-SMA was significantly elevated in the liver of mice treated with CCl_4_, while 3HB could significantly reduce α-SMA levels (Figure 3D & 3G). The level of hydroxyproline in the serum of mice with liver fibrosis could also be significantly reduced by 3HB (Figure 3H), indicating that 3HB can significantly reduce collagen levels in tissues. We also verified the inhibitory effect of 3HB on the expression of α-SMA and collagen in liver tissues, and found that 3HB can significantly reduce the transcriptional level of α-SMA and collagen in the livers of mice with liver fibrosis (Figure 3I & 3J). The above results further validate the therapeutic effect of 3HB on liver fibrosis, and this effect has no negative impact on the body weight and relative liver weight of the mice (Figure 3K & 3L).

### 3HB inhibits the expression of inflammatory factors in mice with liver fibrosis

Previous research has indicated the anti-inflammatory role of 3HB[14], leading us to speculate that its therapeutic effect on liver fibrosis may be associated with this anti-inflammatory capacity. Accordingly, we examined the expression of inflammatory mediators in the serum and liver of mice with liver fibrosis. Our findings revealed that 3HB significantly reduced the elevated levels of IL-1β and TNFα in the serum of mice induced with liver fibrosis by CCl_4_ (Figure 4A-4B). Concurrently in hepatic tissues, 3HB equally diminished the levels of IL-1β and TNFα mRNA in the liver (Figure 4C-4D), demonstrating its potent anti-inflammatory action in mice with liver fibrosis.

**Fig 4.**
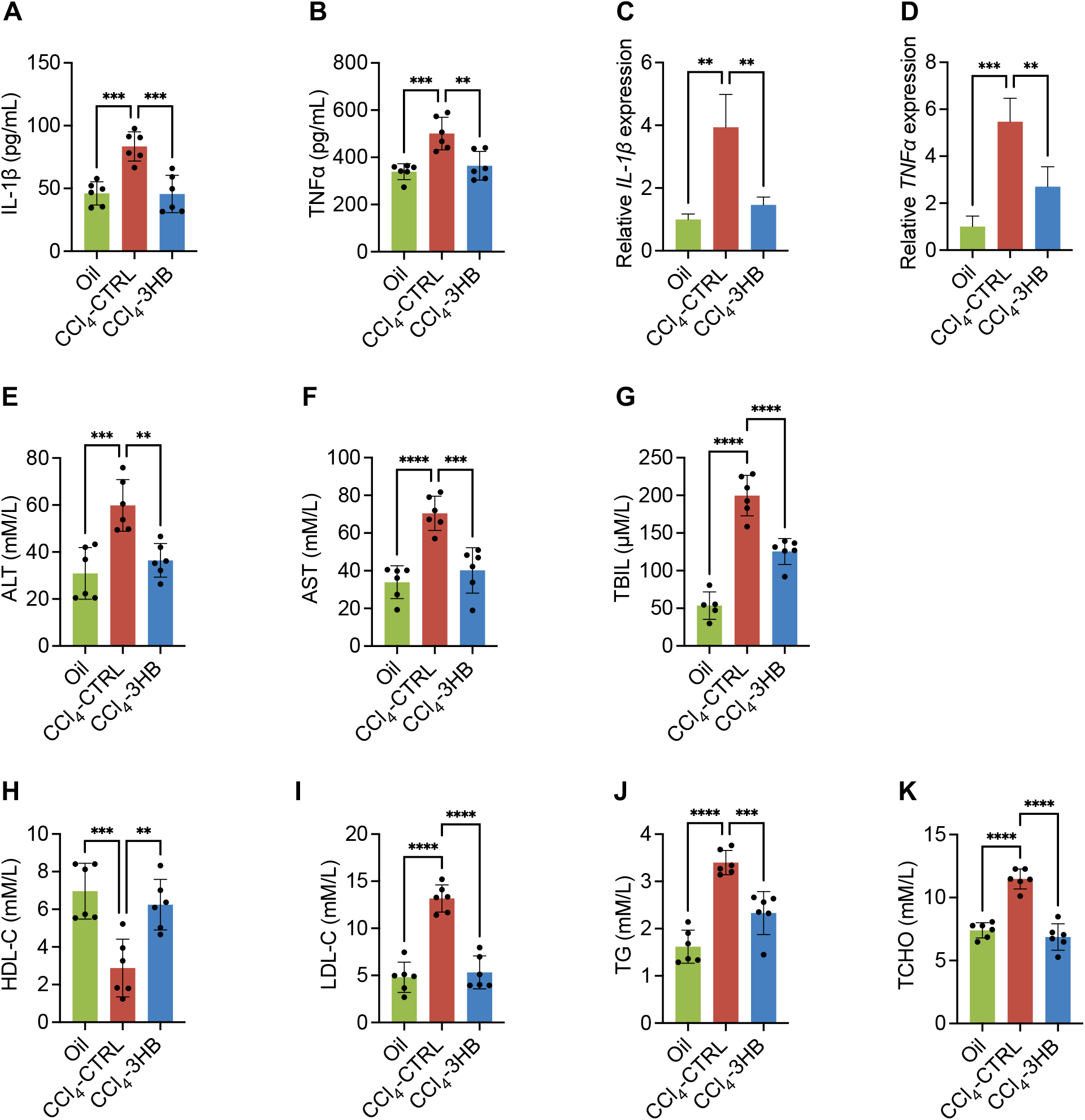
3HB inhibits the expression of inflammatory factors in mice with liver fibrosis. A. Serum IL-1β level. B. Serum TNFα level. C. The mRNA expression levels of IL-1β in liver. D. The mRNA expression levels of TNFα in liver. E. Serum ALT level. F. Serum AST level. G. Serum TBIL level. H Serum HDL-C level. I. Serum LDL-C level. J. Serum TG level. K. Serum TCHO level. Date is presented as means□± □S.E.M. (n□=L6). *p□<L0.05 and **p□<L0.01, ***p□<L0.001, ****p□<L0.0001, compared with CCl_4_-CTRL group.

Similarly, 100 mg/kg of 3HB substantially improved liver function in mice with liver fibrosis and demonstrated significant therapeutic efficacy against hyperlipidemia caused by liver fibrosis (Figure 4E-4K).

### 3HB regulates ECM organization, ROS metabolism, mitochondria fission and p53 pathway

In order to further determine the molecular mechanism of 3HB in mice with liver fibrosis, RNA-sequencing was performed to detect liver gene expression profiles of each group, and cluster map was drawn according to those differentially expressed genes. The heatmap showed that 3HB treatment could significantly alter the gene expression pattern of liver in mice with liver fibrosis (Figure 5A).

**Fig 5.**
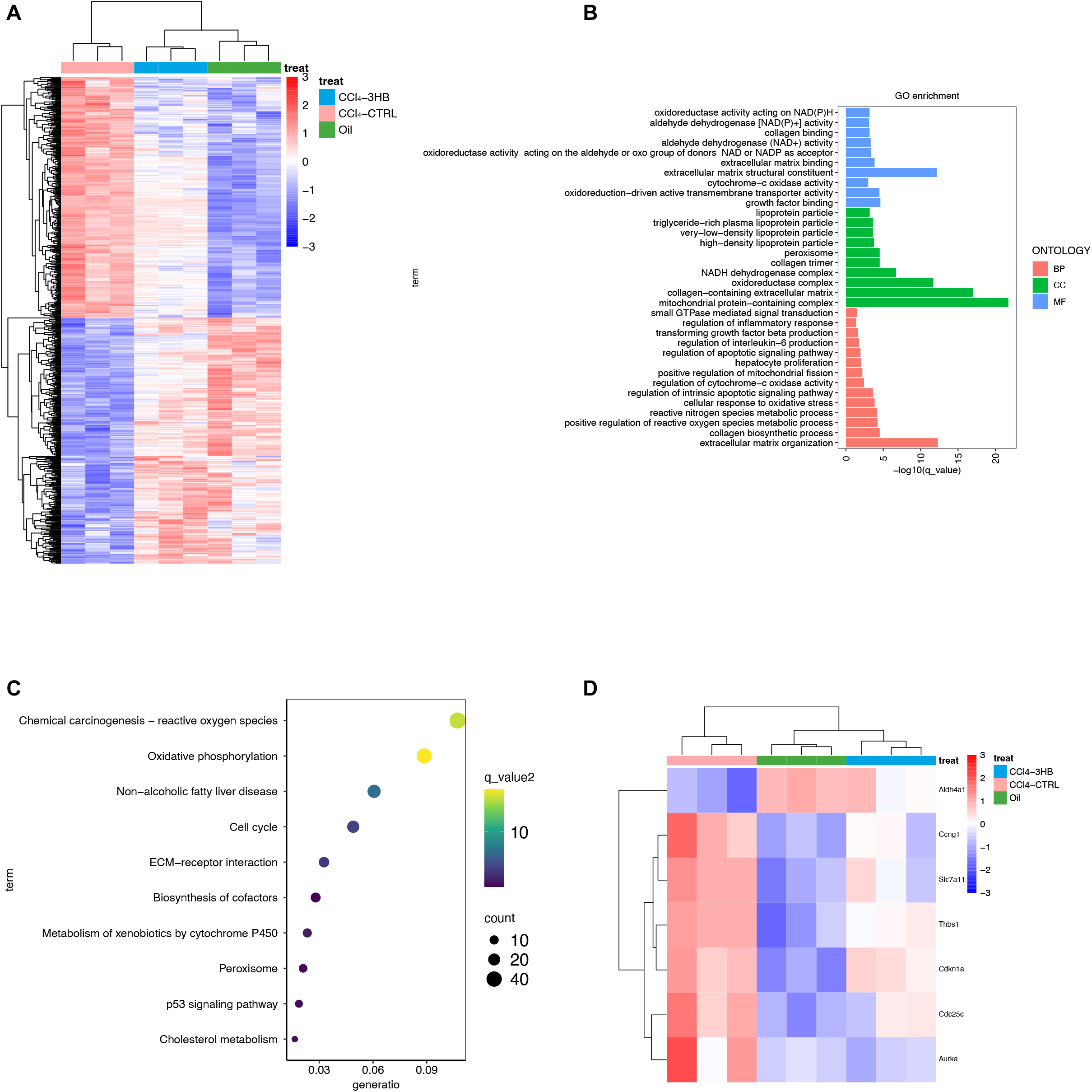
Transcriptome analysis of liver tissue in liver fibrosis treated with 3HB. After 4 weeks of indicated treatment, the liver of mice was collected and studied the differential gene expression analysis (p<0.05) by RNA sequencing. A. Heatmap of differentially expressed genes. Red indicates increased gene expression, while blue indicates decreased gene expression. B. GO enrichment analysis of liver fibrosis mice. The related BP, CC and MF were analyzed. C. KEGG enrichment analysis of pathways down-regulated by 3HB.

Related enriched Gene Ontology (GO) terms were shown in Figure 5B. In terms of biological processes (BP), 3HB was involved in extracellular matrix (ECM) organization, collagen biosynthetic process, reactive oxygen species (ROS) and reactive nitrogen species (RNS) metabolic process, regulation of mitochondrial fission, transforming growth factor beta (TGF-β) production and several processes associated with inflammation in liver. Cellular component (CC) and molecular function (MF) enrichments also indicated that 3HB treatment was closely related to

ECM construction, collagen production, oxidative phosphorylation and mitochondrial function in liver of mice with liver fibrosis. The KEGG enrichment results showed that the regulation of oxidative phosphorylation, ROS, ECM-receptor interaction, p53 and some other signaling pathways were affected by 3HB administration (Figure. 5C). P53 is involved and required in the anti-fibrosis process [19]. A total of 7 reported p53 regulated genes were found out from all differentially expressed genes, and 3HB treatment affected the expression of p53 controlling genes (Figure 5D).

In summary, 3HB may alleviate CCl_4_-induced liver fibrosis by regulating ECM organization, ROS metabolism, mitochondria fission and p53 pathway.

## DISCUSSION

The CCl_4_-induced mouse model of liver fibrosis has been widely used to simulate human liver fibrosis[20]. Our study suggests that 3HB treatment has beneficial effects on CCl_4_-induced liver fibrosis in mice. 3HB can reduce the infiltration of inflammatory cells and the release of pro-inflammatory cytokines induced by CCl_4_, improve liver function in CCl_4_-induced mice, and alleviate the dyslipidemia caused by liver fibrosis. These findings indicate that 3HB may be a potential therapeutic option for the treatment of liver fibrosis.

Previous reports have shown that 3HB can improve alcoholic liver injury and alleviate symptoms of non-alcoholic fatty liver disease (NAFLD) in type 2 diabetic mice, demonstrating its protective effects on the liver[16, 17]. Liver fibrosis is a common pathological consequence of both alcoholic liver injury and NAFLD[21]. Chronic alcohol consumption can lead to the development of alcoholic liver disease, which in its severe form, can progress to liver fibrosis[22]. Similarly, NAFLD, characterized by excessive fat accumulation in the liver, can also advance to liver fibrosis if not appropriately managed[23]. This progression is typically associated with chronic inflammation and the activation of hepatic stellate cells, leading to excessive deposition of extracellular matrix proteins and the development of fibrosis[21]. Therefore, in this study, we explored the role of 3HB in liver fibrosis and found that 3HB significantly inhibited the protein and mRNA levels of collagen and α-SMA in the liver. α-SMA is one of the marker genes of hepatic stellate cell activation[21], and these data suggest that 3HB may inhibit liver fibrosis by suppressing the activation of hepatic stellate cells. Due to reports showing that HSCs can still recover to an inactive state (although not completely quiescent) after activating myofibroblasts, this might be the key factor why 3HB shows such significant effect [13]. Moreover, liver injury leads to increased infiltration of inflammatory cells and release of pro-inflammatory cytokines, which are the main inducers of liver fibrosis[24]. 3HB can significantly reduce the levels of liver damage markers ALT, AST, and TBIL, alleviate CCl_4_-induced liver injury, and protect normal liver function. Previous studies have shown that 3HB has anti-inflammatory effects in animal models of colonic inflammation, atherosclerosis, alcoholic liver injury and depression[17, 25-28]. Our experimental results suggest that 3HB can reduce the infiltration of inflammatory cells in CCl_4_-induced mouse liver and inhibit the expression of pro-inflammatory cytokines IL-1β and TNFα. Considering that these cytokines are inducers of fibrosis, we speculate that 3HB may partially inhibit liver fibrosis by suppressing inflammatory signaling.

To further explore the mechanism of 3HB in treatment of CCl_4_-induced liver fibrosis, we performed transcriptome sequencing analysis to investigate the biological processes and signaling pathways that changed in liver fibrosis tissue after 3HB treatment.

Consistent with the phenotype, 3HB can regulate ECM organization in the liver and regulate collagen formation (Figure 5B). Both GO and KEGG enrichment suggested that 3HB can regulate ROS metabolism (Figure 5B-5C), providing insights for probing the specific anti-fibrosis mechanism of 3HB [29]. Mitochondrial dysfunction is associated with NAFLD and liver fibrosis, and a ketogenic diet can reduce liver injury by restoring mitochondrial function [30, 31]. 3HB is the most significantly elevated molecule during the ketogenic diet process [32]. GO enrichment showed that 3HB regulated the process of mitochondrial fission in the liver (Figure 5B). Therefore, the alleviating effect of 3HB on CCl_4_-induced liver fibrosis may be achieved through the restoring normal function of mouse hepatocyte mitochondria. P53 is involved and required in the anti-fibrosis process [19]. We found that 3HB also regulated the p53 signaling pathway in the liver of mice with liver fibrosis, and affected the expression of p53-related genes. Therefore, exploring the impact of 3HB on the p53 signaling pathway is also one of the options for uncovering the specific molecular mechanisms of 3HB’s anti-fibrosis effect.

## CONCLUSION

In summary, we have found that 3HB can alleviate CCl_4_-induced liver injury and inflammation in mice, slow down the onset of fibrosis, reduce the accumulation of collagen protein and the level of pro-fibrotic gene mRNA, and suppress the expression of inflammatory factors. Our data suggest that 3HB may exert its anti-fibrotic effects through several ways, such as reducing oxidative stress, improving mitochondrial function and p53 signaling pathways, positioning it as a potential option for the treatment or adjunctive therapy of liver fibrosis.

## MATERIALS AND METHODS

### Animal treatments

8-week-old male C57BL/6 mice were purchased from Vital River Laboratory Animal Technology Co, Beijing, China. All mice were housed in a specific pathogen-free animal facility in Tsinghua animal house, with a 12 h light and 12 h dark cycle, with one-week adaptive period before official experiments. All animal procedures were approved by the Institutional Animal Care and Use Ethic Committee of Tsinghua University (NO. 23-CGQ1).

The mice were randomly divided into five groups as following: oil-injected control group, CCl4-injected group, and three 3HB gavage groups with doses of 50, 100 and 200mg/kg. 5ml/kg Oil and CCl4 injected mice were given intraperitoneally pure olive oil and 10% CCl4 (diluted with olive oil) twice a week for two months continuously; 3HB, dissolved in sterile water, was given to mice by gavage every day for three months, with one month of pretreatment before injection. At the end of the animal experiment, all mice were fasted overnight and anesthetized using avertin. Liver tissue and cardiac blood was collected after euthanasia.

### Histology analysis

Collected liver tissues were fixed in 4% paraformaldehyde, embedded in paraffin, cut at 5 μm and stained with Hematoxylin and eosin staining (H&E), Masson trichrome and Sirius red staining according to standard procedures. The stained sections were visualized using a microscopy (Vectra Polaris).

### Biochemical measurements

The blood samples collected from the heart of mice were centrifuged at 4 □, 5000×g for 5 minutes and the supernatants were separated. Indicators for liver function (ALT, AST, TBIL) were detected through corresponding kits; inflammatory factors (IL-1β and TNFα) were detected through corresponding ELISA kits; Serum high density lipoprotein cholesterol (HDL-C), low density lipoprotein cholesterol (LDL-C), triglyceride (TG), total cholesterol (TC) and hydroxyproline were all measured using direct microplate method via kits.

### Isolation of RNA and analysis by real-time quantitative PCR

Total RNA of liver tissues was extracted with TRIzol reagent and the cDNAs were generated by reverse transcription kit. Real-Time Quantitative PCR (qPCR) was performed with the standard protocols in ABI 7500 Fast Real-Time PCR System (Applied Biosystems™ 7500, ThermoFisher, USA) using SYBR Green Master Mix. GAPDH gene expression was used to normalize the expression of mRNA for genes of interest. Primers used for qPCR are listed (Table 1).

**Table 1.**
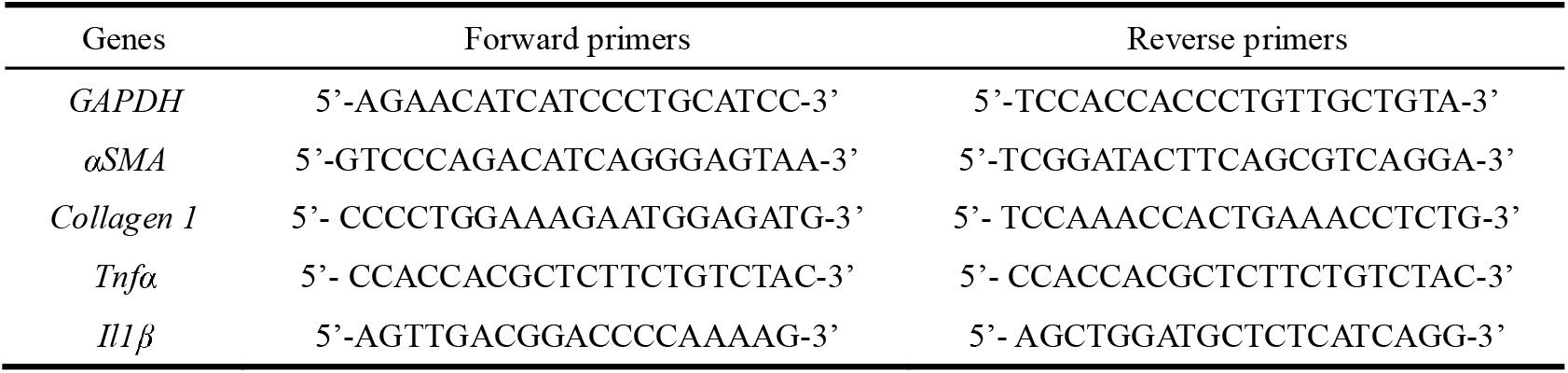
Primers of qPCR.

### Transcriptome sequencing and analysis

Total RNA of adipose tissue was collected to analyze the transcriptome, with a cDNA library constructed using qualified adipose tissue RNA samples. The transcriptome sequencing processes of adipose tissue were performed by Novogene (Beijing, China) using Illumina HiSeqTM 4000 sequencer. The adaptors of paired-end reads were trimmed and quality control checks were carried out using trim-galore (v.0.6.0). Reads were aligned to mouse genome reference (GRCm38.p6) from GENCODE using STAR (v.2.7.3a). FeatureCounts (v.1.6.3) counted reads stored in BAM format to exon sites of genes included in GTF files from GENCODE. Differential gene expression analysis was conducted using raw counts as input using R (v.3.5.1) package DESeq2 (v.1.22.2). Kyoto Encyclopedia of Genes and Genomes (KEGG) analysis and Gene ontology (GO) enrichment analysis of differential genes were performed by R (v.3.5.1) package clusterProfiler (v3.10.1).

### Quantification and statistical analysis

Results are presented as the mean ± standard deviation (SD). The statistical differences between two groups were analyzed using the one-way ANOVA by GraphPad Prism software 9. The data were considered significant difference as the *p* value<0.05.

## DATA AVAILABILITY

All data needed to evaluate the conclusions in the paper are present in the paper. Source transcriptomic data are available through https://www.ncbi.nlm.nih.gov/bioproject/PRJNA1065942. Accession number: PRJNA1065942.

## ACKNOWLEDEMENTS

This research was financially supported by the National Natural Science Foundation of China (Grant No. 31870859) and Tsinghua Chunfeng Foundation.

## ABBREVIATIONS

ALT: Alanine aminotransferase
AST: Aspartate transaminase
α-SMA: Alpha smooth muscle actin
BDH1: 3-Hydroxybutyrate dehydrogenase
1,3-BDO: 1,3-Butanediol
BP: Biological processes
CC: Cellular component
CCl_4_: Carbon tetrachloride
ECM: Extracellular matrix
GO: Gene ontology
HCC: Hepatocellular carcinoma
HDL-C: High-density lipoprotein cholesterol
3HB: 3-Hydroxybutyrate
H&E: Hematoxylin and eosin staining
IL-1β: Interleukin 1β
KEGG: Kyoto encyclopedia of genes and genomes
LDL-C: Low-density lipoprotein cholesterol
MF: Molecular function
NAFLD: Non-alcoholic fatty liver disease
NASH: Nonalcoholic steatohepatitis
NLRP3: NOD-like receptor family: pyrin domain: containing 3
ROS: Reactive oxygen species
TBIL: Total bilirubin
TG: Total triglycerides
TCHO: Total cholesterol
TNFα: Tumor necrosis factor α
TGF-β: Transforming growth factor β

## AUTHOR CONTRIBUTIONS

G.-Q.C. and Y.Z. supervised this study. Y.Z. and X.L. designed and performed the overall research, and wrote the manuscript. Y.W., M.L, Z.G., J.Z. and F.W. provided support with experimental techniques and revised the manuscript. All authors have read and approved the article.

